# Anp32e protects against accumulation of H2A.Z at Sox motif containing promoters during zebrafish gastrulation

**DOI:** 10.1101/2023.12.18.572196

**Authors:** Fabian N. Halblander, Fanju W. Meng, Patrick J. Murphy

**Affiliations:** Department of Biomedical Genetics, University of Rochester Medical Center, Rochester NY, 14642, USA

**Keywords:** H2A.Z, Anp32e, chromatin accessibility, Sox, neural crest, embryonic development

## Abstract

Epigenetic regulation of chromatin states is crucial for proper gene expression programs and progression during development, but precise mechanisms by which epigenetic factors influence differentiation remain poorly understood. Here we find that the histone variant H2A.Z accumulates at Sox motif-containing promoters during zebrafish gastrulation while neighboring genes become transcriptionally active. These changes coincide with reduced expression of *anp32e*, the H2A.Z histone removal chaperone, suggesting that loss of Anp32e may lead to increases in H2A.Z during differentiation. Remarkably, genetic removal of Anp32e in embryos leads to H2A.Z accumulation prior to gastrulation, and precocious developmental transcription of Sox motif associated genes. Altogether, our results provide compelling evidence for a mechanism in which Anp32e restricts H2A.Z accumulation at Sox motif-containing promoters, and subsequent down-regulation of Anp32e enables temporal up-regulation of Sox motif associated genes.

**Highlights:**

- An early-developmental time course of zebrafish chromatin accessibility is achieved using an integrated UMAP analysis of datasets from two separate published studies.
- CUT&Tag sequencing is used to characterize the genomic localization for the histone chaperone ANP32E.
- Changes in Anp32e enrichment coincide with opposing changes in H2A.Z enrichment during zebrafish gastrulation.
- Precociously expressed genes in embryos lacking Anp32e are disproportionately Sox-marked and may represent H2A.Z-mediated developmental accelerations.

## Introduction

Epigenetic marks on chromatin, including post-translational modifications to canonical histones and the incorporation of histone variants, can play a key role in regulating gene expression and transcription factor (TF) function^1^. Previous studies have shown that cell state transitions during development are often accompanied by major epigenetic changes^2,3,4^, but the means by which these changes influence discrete biological pathways remain only partially understood.

The histone variant H2A.Z is highly conserved across organisms and is essential for early embryogenesis in numerous animals^5,6^. H2A.Z histones are deposited primarily at gene regulatory regions within the genome, including at promoters surrounding the transcription start site (TSS) and enhancers^7,8^. Nucleosomes containing H2A.Z have been described to be more labile than canonical nucleosomes^9,10^, potentially influencing how transcription factors gain access to DNA binding sites or the transcription levels that occurs for a given gene^11^. Several studies, including ours, have shown that depletion of the H2A.Z removal chaperone, ANP32E, leads to H2A.Z accumulation at promoters, which is accompanied by increased chromatin accessibility and amplified gene expression levels^12–13^. While these studies provide compelling molecular mechanisms by which precise control of H2A.Z could regulate cellular gene expression, it remains unknown whether this mode of regulation contributes to specific processes involved in embryonic development.

In zebrafish, inheritance of H2A.Z-containing placeholder nucleosomes prepares the embryo for zygotic genome activation^14^, but the function of these nucleosomes following the initial cleavage phase has not been explored. Interestingly, separate studies have implicated H2A.Z to be particularly important for the development of neural crest derived tissues^15,16^, but considering the ubiquitous nature of H2A.Z expression in animals, and its necessity during early development^5^, it remains poorly understood how tissue-level specificity is attained. In prior studies, we identified human *ANP32E* expression patterns to be anti-correlated with breast cancer disease progression, and we found H2A.Z localization to be correlated with high levels of chromatin accessibility at several TF binding sites, including motifs for SOX factor and FOX factor binding^17^. One compelling possibility is that function of these types of TFs is particularly impacted by H2A.Z localization, and in this manner, changes in H2A.Z localization could specifically influence the expression of genes regulated by these factors. Numerous SOX and FOX family TFs are known to regulate specific aspects of development, including the formation of multiple adult stem cell populations and neural crest cells^18–20^. Whether these factors are influenced by ANP32E or H2A.Z in the context of development remains unknown.

Here, we used zebrafish embryos to Investigate the role of H2A.Z during the developmental transition from pluripotency through gastrulation and early differentiation. We integrated and reanalyzed a time course of publicly available chromatin accessibly (ATAC-seq), RNA-Seq, and ChIP-Seq datasets ^2,21^, along with our newly acquired genomic CUT&Tag profiles for H2A.Z and Anp32e. We find that Anp32e resides primarily at Sox motif containing gene promoters in early embryos and Anp32e binding decreases as embryos transition through gastrulation. We also find that H2A.Z enrichment increases at these loci during this same developmental period.

Through subsequent measurements of H2A.Z localization and gene expression changes in embryos lacking Anp32e, we are able to formulate a model in which Anp32e binding protects Sox motif containing promoters against accumulation of H2A.Z at the early stages of zebrafish development.

## Results

### H2A.Z accumulates at putative regulatory elements during early differentiation

To investigate epigenetic patterning during the initial phase of zebrafish differentiation, we analyzed enrichment for histone modifications H3K4me1, H3K4me3, and H3K27ac using publicly available datasets^22^. Datasets for each histone modification were collected in blastula embryos (dome stage) and gastrula embryos (80% epiboly)^23^. For each dataset, we then defined genomic locations where enrichment was strongest, termed ‘peaks’ (see methods), and further identified developmental stage-specific enrichment, based on whether peaks overlapped across timepoints (Fig. 1A & 1B). This strategy enables us to identify loci in which levels of H3K4me1 and H3K4me3 increased during differentiation (Fig. S1A), referred to as ‘gastrulation-specific’. As anticipated, developmental accumulation of either H3K4me1 or H3K4me3 associated with higher levels of H2A.Z, H3K27ac, and chromatin accessibility (ATAC-seq)^2^ (Fig. 1C-D). Gene ontology analysis revealed that genes located in proximity to these sites tended to be involved in embryonic development and metabolism (Fig. S1D), and interestingly, binding motifs for several transcription factors involved in neural crest development were found to be enriched (Fig. 1E), including Sox family, Fox family, and Dlx factors. These results are consistent with our hypothesis that accumulation of H2A.Z may contribute to stage-specific developmental epigenetic regulation.

**Figure 1:**
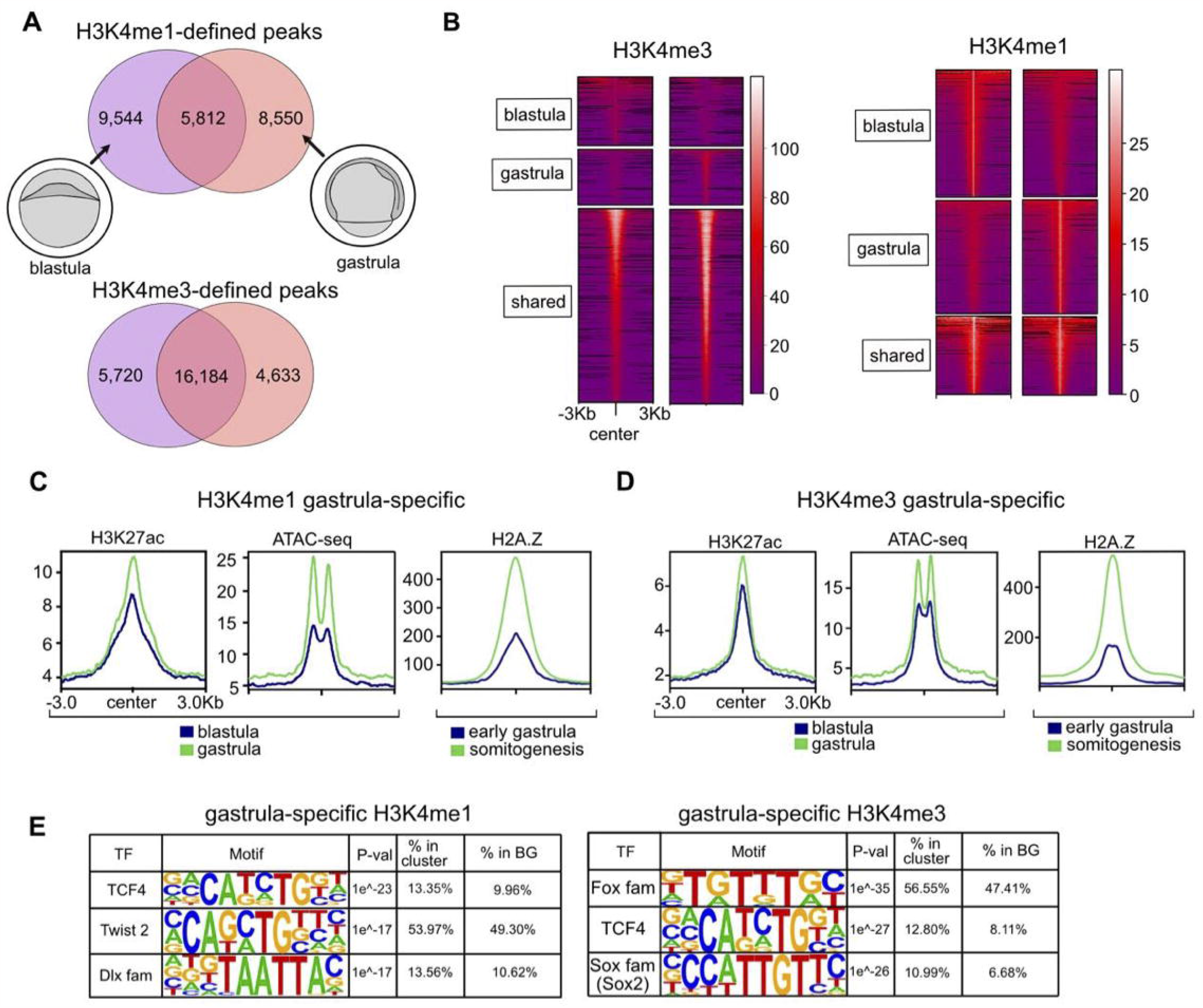
H2A.Z accumulates at gastrulation-specific putative regulatory elements. A. H3K4me1 and H3K4me3 peaks (ChIP-seq) are partitioned into stage-specific classes. Venn diagram depicts the number of peaks that are blastula-specific (purple), gastrula-specific (pink), or maintained in transition (blend of colors). B. Heatmaps H3K4me3 and H3K4me1 at blastula-specific, gastrula-specific, and shared peaks. In either H3K4me1 or H3K4me3, Blastula-specific regions decrease, gastrulaspecific regions increase, and shared regions maintain. C. Genomic averages of H3K27ac (ChIP-seq), chromatin accessibility (ATAC-seq), and H2A.Z (CUT&Tag) at H3K4me1 gastrula-specific regions. D. Analogous to panel C at H3K4me3 gastrula-specific regions. E. Selection of transcription factor motifs enriched in gastrula-specific H3K4me1 and H3K4me3 regions. Enrichment is found for TCF, Twist, Dlx, Fox, and Sox factors between categories.

### Increases in chromatin accessibility occur during gastrulation over Sox motif containing promoters

We next assessed whether developmental chromatin changes coincide with epigenetic changes, and utilized a series of published datasets to survey a developmental window spanning zebrafish gastrulation (utilizing publicly available bulk ATAC-Seq data^2^). For comparison, we also included ATAC-seq datasets acquired from purified neural crest cells^21^. Published pre-processed early embryonic RNA-seq data was also collected^24^, allowing us to compare chromatin changes with transcriptional impacts. After rank-normalization we performed UMAP analysis, and the resulting profiles parallel the developmental progression (Fig. 2A – green arrow). We observed a clear separation of datasets between UMAP dimensions 1 and 2, revealing there to be a distinction in chromatin states for undifferentiated and differentiated embryos (Fig. 2A – red line). Samples from later developmental stages (including shield, epiboly, and somitogenesis) had negative values for both dimensions, whereas earlier stages (including high, oblong, sphere, and dome) had positive values. In agreement with our UMAP analysis, K-means clustering of rank-normalized chromatin accessibility scores also revealed there to be a clear separation between samples from differentiated and undifferentiated embryos (Fig. S2A).

**Figure 2:**
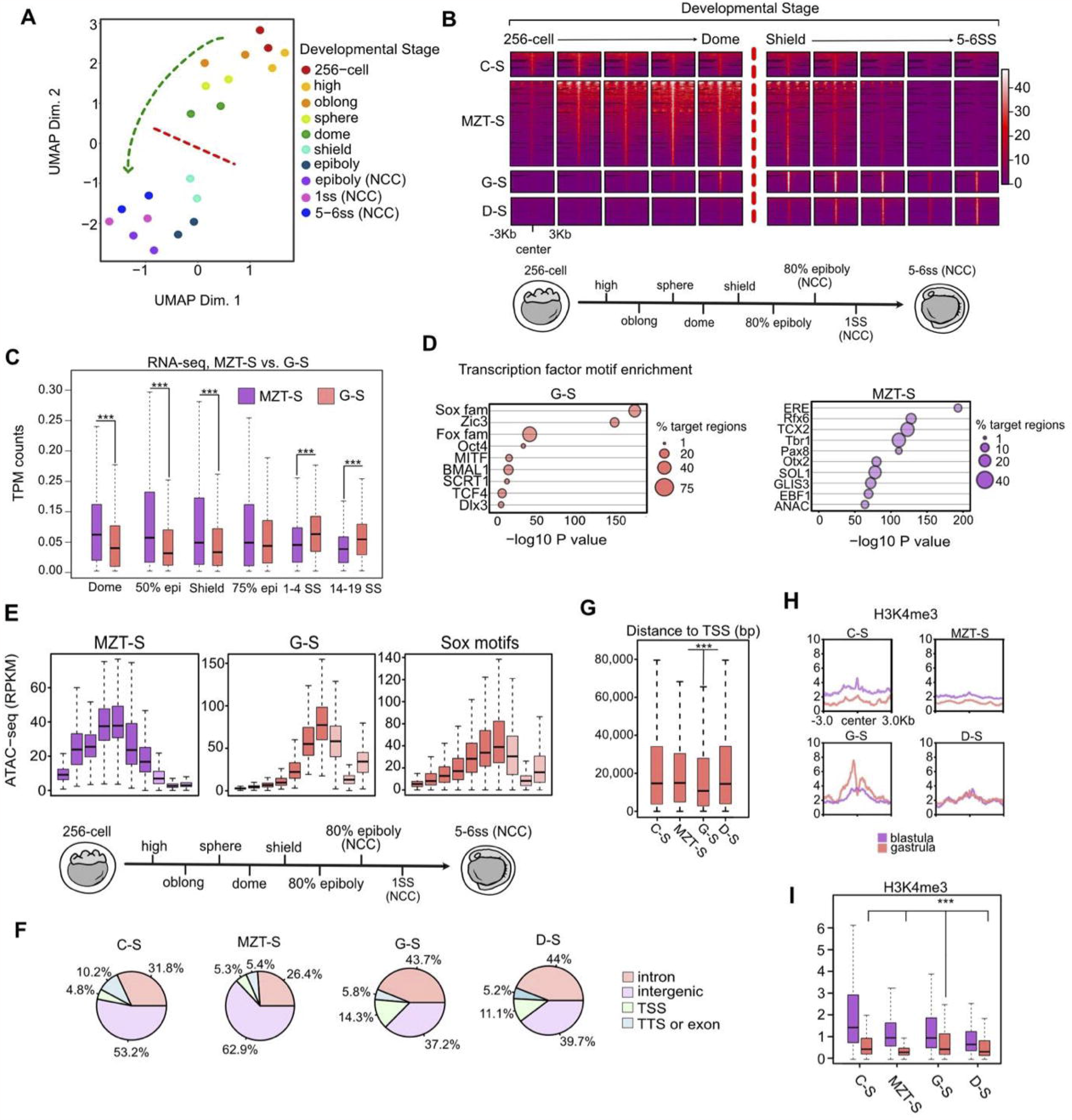
Changes of chromatin accessibility across developmental stages. A. UMAP projection of a rank-normalized chromatin accessibility (ATAC-seq) profile recovers developmental stage of inputted samples. A genome-wide restructuring of chromatin state (motion of the green-arrow) distinguishes cleavage and blastula periods (positive values of dimension 2, above the red-line) from gastrulation and early differentiation periods (negative values of dimension 2, beneath the red-line). Datasets including and following epiboly are from purified Neural Crest Cell (NCC). Others are whole-embryo. B. Feature-to-feature visualization of read-count normalized accessibility data underlying UMAP analysis, classified by accessibility signature. Cleavage-Specific (C-S, n=1,105), MZT-Specific (MZT-S, n=3,873), Gastrulation Specific (G-S, n=969), and Differentiation-Specific (D-S, n=1,247) classes synchronize with different developmental periods. C. TPM-normalized gene-expression (RNA-seq) at genes nearest G-S and MZT-S loci. MZT-S-associated transcription is higher during MZT. G-S-associated gene transcription is increasing from shield stage (gastrulation) and overtakes MZT-S transcription by 1-4SS (neurula). (p< 0.001 for each). D. Transcription factor motif enrichment is displayed for G-S (left) and MZT-S (right). Motif and motif families are sorted by -log10 P-value measuring statistical significance. Motif % enrichment in target regions is visualized. E. Accessibility (ATAC-seq) patterns at Sox family motifs parallel patterns at G-S regions. This trend is contrasted by MZT-S regions. F. Breakdown of cluster gene annotation frequencies, where colors indicate annotation type. C-S and MZT-S regions are dominated by intergenic sequences. G-S and D-S regions are dominated by genic sequences. Of any class, G-S regions have the highest proportion of TSS. G. Absolute distance to TSS (base-pairs) for each cluster is also displayed. G-S regions are closest to TSS (p <0.001). H. Blastula and gastrula H3K4me3 enrichment in each accessibility class. Genomic averages suggest that H3K4me3 increases only at G-S regions. I. Boxplots analogous to profile plots in H. G-S regions have the highest H3K4me3 at gastrula.

We then classified genomic loci based on the developmental stage in which chromatin accessibility reached it maxima. Loci were classified as either cleavage stage specific (most accessible shortly after fertilization), maternal zygotic transition (MZT) specific (most accessible prior to gastrulation), gastrula-specific (most accessible around shield and epiboly stages), or differentiation-specific (most accessible during somite stages or in differentiation cells) (Fig. 2B). As anticipated, chromatin changes during the transition from MZT to gastrulation were associated with likewise developmental changes in gene expression (Fig. 2C). For example, genes nearby MZT-specific loci were more highly expressed during MZT. As gastrulation proceeds, MZT-specific transcription diminishes while gastrula-specific transcription intensifies (Fig. 2C). Comparisons between gene ontology terms (from genes proximal to accessibility sites) indicated that developmental changes in chromatin state might be particularly important for gene pathways associated with neural development, cell migration, and morphogenesis (Fig. S2B). Gastrula-specific accessible sites were also strongly enriched for SOX family transcription factor binding motifs (Fig. 2D), similar to our prior measurements of loci which gain H3K4me3 during gastrulation (Fig. 1E).

Having found that Sox motifs were enriched within gastrula-specific accessible sites, we next investigated whether chromatin accessibly changes occurred across all Sox motifs throughout the genome during early development. Remarkably, changes in chromatin accessibility at Sox motifs were almost identical to the developmental change we observed at gastrula-specific loci (Fig. 2E), with accessibility being low during cleavage phase and then increasing dramatically during gastrulation (in shield and epiboly stage embryos). We also found that gastrula-specific accessible regions contained more genic loci (including TSS and intronic sequences) than MZT-specific regions contained (Fig. 2F), and gastrula-specific regions were in closer proximity to transcription start sites as compared with other classes of accessible loci (Fig. 2G), suggesting that chromatin reprogramming may occur to a greater extent over gene promoters during gastrulation than during other developmental periods. As further support for promoter-specific impacts, we found that levels of H3K4me3 (which predominantly marks promoters^25^) were higher in gastrula stage embryos at gastrula-specific accessible loci, as compared with MZT-specific or differentiation-specific regions (Fig. 2H & 2I). Taken together, these data suggest that chromatin becomes more accessible over gene promoters during zebrafish gastrulation, SOX motifs tend to be enriched at these loci, and up-regulation of proximal gene expressions occurs in association with increased accessibility.

### Loss of Anp32e over Sox motif containing promoters leads to H2A.Z accumulation

Having identified genomic loci possessing clear indicators of chromatin reprogramming during gastrulation (increased accessibility, H3K4me3 enrichment, and increased neighboring gene activation), we then returned to our initial hypothesis, that H2A.Z accumulation during early development helps to support gene activation. In agreement with this hypothesis, we found that developmental accumulation of H2A.Z occurred over a majority of gene promoters during gastrulation (Fig S3A), and similar patterns were observed for gastrula-specific accessible regions located within promoters (Fig 3A & 3B), but not for regions located outside of promoters. Additionally, H2A.Z levels were significantly higher over promoters containing at least one Sox factor binding motif (Fig 3C & 3D), compared with those lacking a motif.

In prior studies, we and others demonstrated that H2A.Z accumulation is restricted by the histone chaperone Anp32e in vertebrates^12–14^. We therefore speculated that the observed H2A.Z accumulation during gastrulation might have been the result of Anp32e loss during these timepoints. Along these lines, we found that *anp32e* gene expression levels decreased nearing the end of gastrulation, unlike expression of *h2afva* or *h2afvb* (Fig 3E). Furthermore, CUT&Tag measurements revealed that high levels of Anp32e occurred over the majority of promoters during early gastrulation (shield stage), but levels were reduced by somitogenesis (5-6SS), encompassing the end of gastrulation (Fig S3B). As anticipated, similar changes in Anp32e levels were observed for gastrula-specific regions classified as promoters, but not for regions classified as non-promoters, which maintained low levels of Anp32e (Fig 3F & 3G). We also found that Anp32e levels were significantly higher over promoters containing a Sox motif, compared with promoters lacking a motif, and Anp32e levels decreased over these promoters during gastrulation (Fig 3H & 3I).

**Figure 3:**
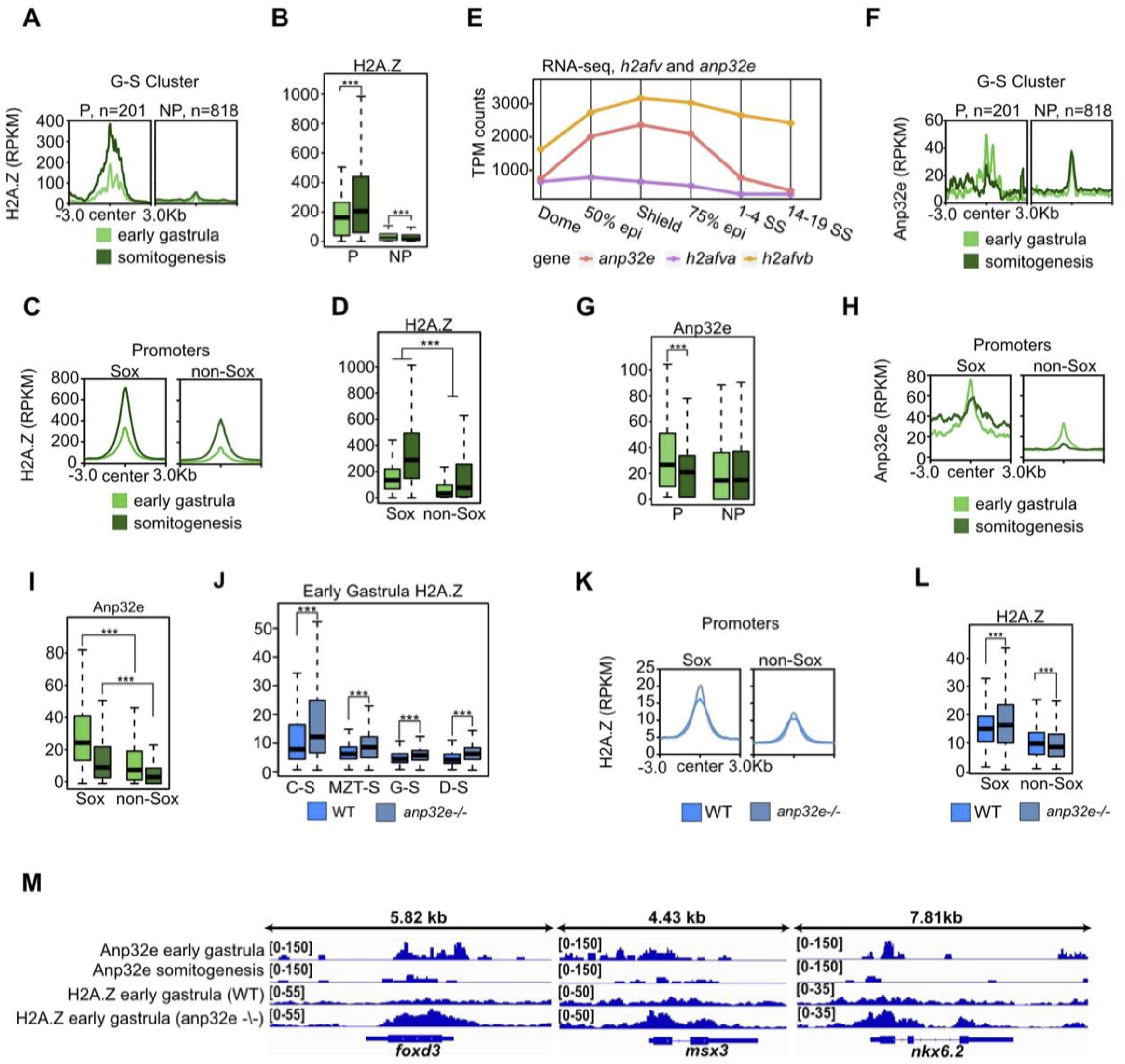
Anp32e loss leads to H2A.Z accumulation at Sox motif containing promoters. A. H2A.Z enrichment averages (CUT&Tag) for early (6hpf) and late (12hpf) gastrula stage embryos is displayed at G-S regions partitioned into promoters (P, n=201) and non-promoters (NP, n=818). B. Boxplots of H2A.Z levels in early and late stage gastrula embryos demonstrate increases at G-S promoters and decreases at G-S non-promoters during gastrulation (p<0.001). C. H2A.Z enrichment averages in early and late stage gastrula embryos at promoters containing Sox motifs (n=6,968, Sox) and promoters not containing Sox motifs (n=25,058, nonSox). D. Boxplots of H2A.Z levels in early and late gastrula embryos demonstrate that H2A.Z is more highly enriched at Sox promoters (p<0.001). E. Gene expression levels (RNA-seq) of *anp32e, h2afva, and h2afvb* from Dome (blastula) to 14-19SS (neurula). *h2afva* and *h2afvb* are increasing from dome (blastula) and are stably expressed after 75% epiboly (gastrulation). Expression of *anp32e* is reduced after 75% epiboly. F. ANP32E enrichment averages in early and late gastrula embryos at G-S regions partitioned into promoters (P) and non-promoters (NP). G. Boxplots of Anp32e levels in early and late gastrula embryos demonstrate decreases at G-S promoters (p<0.001). Anp32e stays the same at G-S non-promoters. H. Early and late gastrula average Anp32e enrichment at Sox and non-Sox promoters. I. Boxplots of early and late gastrula Anp32e levels show higher levels at Sox promoters than non-Sox promoters for each sample (p<0.001). J. Boxplots of H2A.Z levels in WT and anp32e -/- embryos at 6hpf (ChIP-seq). H2A.Z is measured at each of our previously identified accessibility clusters. H2A.Z increases at all regions in anp32e -/- embryos (p<0.001). K. Average enrichment of H2A.Z at Sox and non-Sox promoters in WT and anp32e -/-. L. Boxplots of H2A.Z levels at Sox and non-Sox promoters in WT and anp32e -/- embryos. Sox promoters increase in H2A.Z in KO embryos compared to WT at early gastrul A. Non-Sox promoters decrease. M. Genome browser screenshots of Anp32e and H2A.Z enrichment in WT and anp32e -/- embryos at early gastrul A. Sox promoter genes *foxd3, msx3, and nkx6*.*2* are displayed. At these sites, Anp32e decreases over gastrulation, and excess H2A.Z accumulates in anp32e -/- embryos.

To investigate the extent to which Anp32e might restrict H2A.Z accumulation, we next analyzed H2A.Z enrichment in embryos lacking Anp32e (at shield stage). Loss of Anp32e led to an overall increase in H2A.Z levels over promoters (Fig. S3C), as well as across all previously defined classes of chromatin accessibility (Fig. 3J). Despite only a modest change in average levels (Fig. 3K), the overall distribution measurements indicated that H2A.Z levels increased specifically for promoters containing a Sox motif (Fig. 3L). Taken together, these results suggest that Anp32e functions to restrict H2A.Z accumulation during the earliest stages of zebrafish development, spanning cleavage stage, MZT, and blastula, and during subsequent stages of gastrulation, decreases in Anp32e levels allow H2A.Z to accumulate over gene promoters, with sites containing Sox motifs most affected.

### Anp32e loss leads to precocious developmental transcription of Sox-motif associated genes

Prior studies have reported that increases in H2A.Z levels over promoters corresponds with increased gene expression^5,11^, leading us to speculate that presence of Anp32e may help to dampen transcription by restricting H2A.Z accumulation^12,13,26^. To investigate this possibility, we reanalyzed published RNA-Seq data and compared expression patterns from Anp32e null embryos with expression patterns at other developmental timepoints, both before and after the transition from blastula to gastrula. To minimize expression differences caused by batch-to-batch variation, we rank normalized gene expression levels and performed Principal Components (PC) Analysis. Strikingly, the vast majority of identified sample-to-sample variation occurred along PC2 (ranking from -0.4 to +0.4) as compared with PC1 (ranging from -0.37 to - 0.28) (Fig. 4A), and datasets aligned vertically along PC2 according to developmental stage. Samples from embryos prior to MZT had the most positive PC2 values (+0.4) whereas samples from gastrula embryos had the most negative PC2 values (-0.4) (Fig. 4A). Interestingly, samples from blastula stage embryos lacking Anp32e had slightly lower values for PC2 (-0.2) compared with WT blastula stage embryos (-0.1), indicative of a modest developmental advancement in transcriptional patterns.

**Figure 4:**
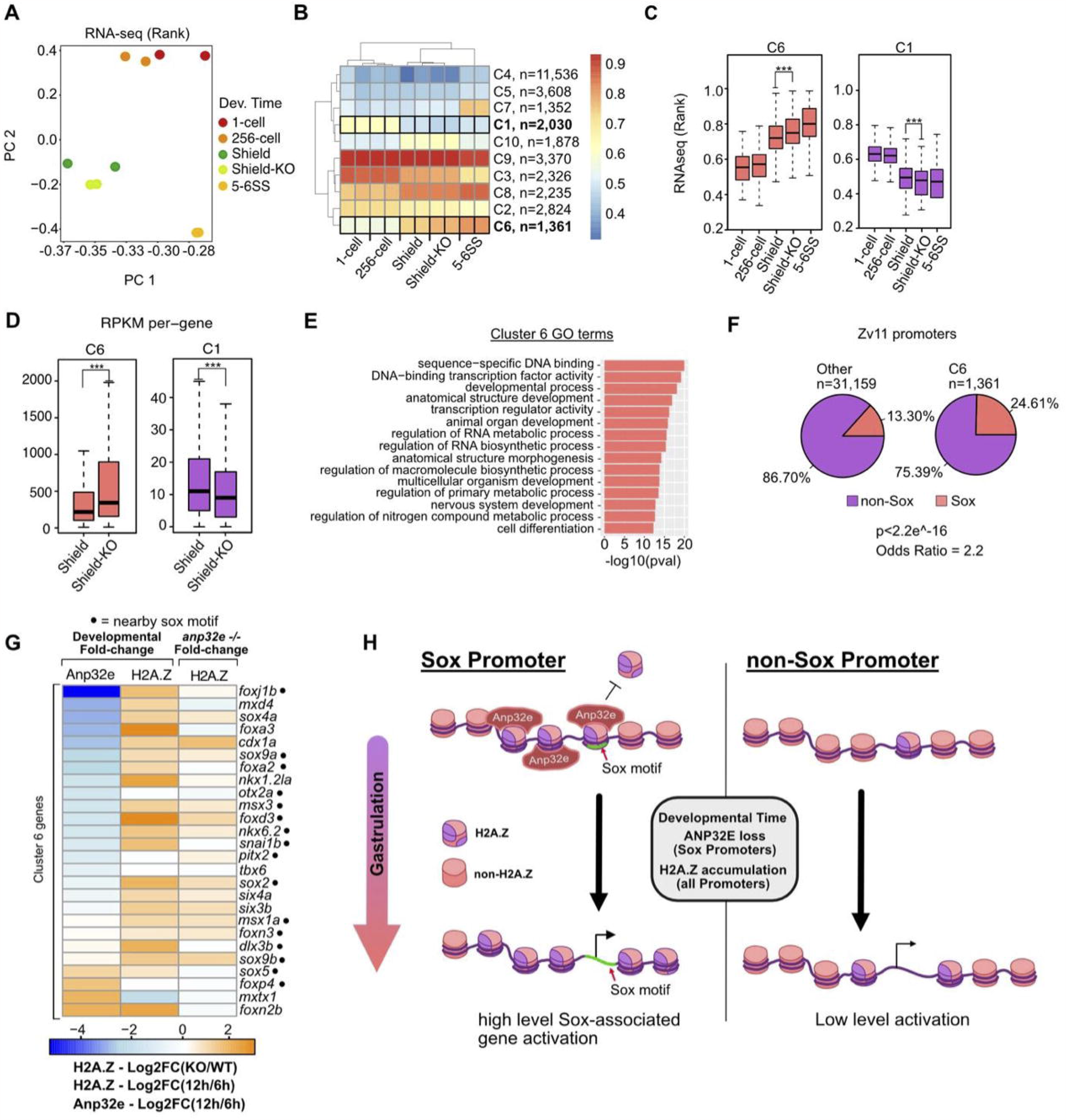
Anp32e loss leads to precocious developmental transcription of Sox-motif associated genes. A. PCA of a time course of rank normalized WT RNA-seq data for 1-cell, 256-cell, Shield, and 5-6SS, as well as anp32e -/- at shield stage (Shield-KO). B. Heatmap of data underlying PCA analysis (k-means=10). Columns are grouped by sample. Rows are grouped by k-means clustering. Shield-KO for C1 and C6 show the clearest differences in expression. C. Boxplots of rank normalized RNA-seq of identified clusters. C6 Shield-KO increase slightly to approach the expression levels of 5-6SS embryos, and C1 Shield-KO decrease. One-tailed t-tests were used to establish that G-KO expression levels increase or decrease compared to G (p<0.001). D. RPKM of genes within C1 or C6. Trends identified in clusters are maintained in the absence of rank-normalization (p<0.001). E. A list of C6 GO terms. Redundant GO terms were removed and a cutoff of >=5 was set for -log10(p-value). F. Fraction of promoters containing one or more Sox motif in C6 versus all other genes. Fischer’s exact test and odds-ratio quantify difference of Sox and non-Sox fractions across gene set promoters. G. A selection of C6 genes. Fold-change of H2A.Z (12hpf/6hpf), Anp32e (12hpf/6hpf), and H2A.Z (KO/WT) are quantified at corresponding promoters. Promoters are marked for the presence of an embedded or adjacent Sox motif. H. A model for Sox-specific developmental regulation of H2A.Z by Anp32e is depicted. High Anp32e levels at the beginning of gastrulation prevent H2A.Z accumulation, thus maintaining transcriptional silencing at sox-motif-containing promoters, and developmental downregulation of Anp32e leads to H2A.Z accumulation, and neighboring gene upregulation

We next performed unbiased K-means clustering analysis, in order to parse genes into clusters based on differences in expression patterns. Similar to our PC analysis, samples organized according to their developmental stage, and several discrete clusters of expression changes emerged (Fig. 4B). Initial comparison of early gastrula stage (6 hours post fertilization) WT and Anp32e deficient embryos revealed there to be little differences between sample types, aligning with prior reports of modest phenotypic differences at this timepoint^14^. We did however detect differences in expression within two clusters – C1, which exhibited reduced expression as compared with WT, and C6 which exhibited increased expression (Fig. 4C & 4D). GO analysis revealed there to be no significant ontology terms for genes within C1, but for C6 several developmentally important ontology terms emerged, including Anatomical Structure Development, Nervous System Development, and Cell Differentiation (Fig. 4E). Based on the observed specificity for H2A.Z accumulation at Sox-motif-containing regions noted previously (Figs 2 and 3), we next assessed enrichment for Sox motifs. Analogous to our H2A.Z measurements, Sox motifs were found to be enriched within cluster C6 gene promoters (p<10^-16,^ odds ratio = 2.2), and not within C1 gene promoters (Fig. 4F). Further comparison of selected genes indicated that numerous Sox motif containing genes within C6 gained H2A.Z and lost Anp32e during gastrulation, and these same genes precociously gain H2A.Z in blastula stage embryos lacking Anp32e (Fig. 4G). These data provide compelling evidence for a common regulatory mechanism in which Anp32e binding at the promoters of Sox motif containing genes protects against H2A.Z accumulation in blastula stage embryos, and subsequent developmental loss of Anp32e (during gastrulation) leads to a buildup of H2A.Z and transcriptional upregulation (Fig. 4H).

## Discussion

Proper control of chromatin state is crucial for regulation of gene expression programs, which is essential for embryonic development and subsequent lineage diversification. In this work, we demonstrate that Anp32e binding at Sox motif containing promoters prevents H2A.Z accumulation at blastula stage. We find that *anp32e* expression decreases as embryos further develop into gastrula stage, concurrent with H2A.Z accumulation at promoters, increases in chromatin accessibility, and activation of genes associated with early development. Strikingly, Anp32e loss leads to H2A.Z accumulation and precocious developmental transcription of Sox-motif associated genes, further highlighting the importance of Anp32e and H2A.Z in regulating developmental gene expression programs. Thus, precise control of level and localization of Anp32e and H2A.Z offer an intriguing mode for the governance of developmental programs in early embryos.

Here we show that Sox motifs are enriched within the genomic loci where chromatin accessibility and H2A.Z enrichment changes occur, suggesting that H2A.Z localization may be particularly important for the function of certain transcription factors or the development of certain cell types. In support of this possibility, recent studies indicate that mammalian SOX9 alters chromatin landscape by binding to regions containing H2A.Z, potentially facilitating chromatin accessibility and activating histone modifications during development^27^. Interestingly, Sox factors are well established to be key players in the speciation and differentiation of a highly specialize vertebrate cell type, termed neural crest cells (NCCs). These cells acquire multipotency after gastrulation during development, allowing them to differentiate into a wide range of embryonic cell types, including cardiomyocytes, cartilages, melanocytes, and neurons^28^. In the context of our study, one interesting possibility is that NCC development might be particularly sensitive to H2A.Z changes, potentially via disruption of Sox factors. Indeed, H2A.Z has been previously shown to be important for the development of NC-derived melanocytes in zebrafish^16^, and mutations in the H2A.Z installation machinery^16^, or disruption of H2A.Z interacting factors PWWP2A and HMG20A^29,30^, cause strong disruption of in NCC-derived tissues, including craniofacial cartilages in *Xenopus Leavis*. In the context of our results, these studies suggest that specific transcription factors are critically important for NCC development, and H2A.Z enrichment might be particularly important for the function of these factors. Additionally, it is quite possible that acquisition of multipotency in NCCs requires extensive chromatin remodeling and this process depends on extensive H2A.Z changes. Evidently, future studies are necessary to examine the genomic dynamics of H2A.Z during NCC development, and its function with respect to Sox factors.

Often times, developmentally important molecular and genetic mechanisms can become reactivated or dysregulated during disease formation. For examples, Sox factors, which have been found to be key regulators of adult stem cell populations, can be hijacked to promote tumor formation during carcinogenesis^18^. Interestingly, recent studies have implicated a role for Anp32e and H2A.Z in multiple types of cancer, including uterine leiomyoma and breast cancer ^17,31,32^. In this context, the model that we propose here, whereby Anp32e and H2A.Z impact transcription factor activity and cell state transition during embryogenesis, may also influence these cancers. Indeed, we previously demonstrated that chromatin accessibility at FOX motifs is associated with breast cancer progression, and is negatively associated with ANP32E expression levels, suggesting that ANP32E might restrict chromatin changes at FOX motifs during breast tumorigenesis^17^ – perhaps in a manner similar to the mechanism we describe for Sox motif containing genes during zebrafish development. In this light, the outcomes of this current study may provide novel insights into mechanisms controlling cellular programming during disease onset and/or progression.

### Key Resources Table

| Reagent or resource | Source | Identifier |
| --- | --- | --- |
| **Antibodies** |  |  |
| Rabbit polyclonal antibody against H2A.Z | Active Motif | Cat. No. 39113 |
| Rabbit polyclonal antibody against Anp32e | ThermoFisher Scientific | Cat. No. PA5-42860 |
| **Critical Commercial Assays** |  |  |
| High-Fidelity 2X PCR Master Mix | NEB | M0541S |
| **Deposited Data** |  |  |
| Fig.1, Fig.2 ChIP-seq | Bogdanovic 2012 | GEO: GSE32483 |
| Fig.1, Fig.2, Fig.3, ATAC-seq (bulk) | Palfy 2020 | GEO: GSE130944 |
| Fig.2, Fig.3, ATAC-seq (NCC) | Lukoseviciute 2018 | GEO: GSE106676 |
| Fig.2 RNA-seq | White 2017 | ENA: PRJEB7244, PRJEB12296, PRJEB12982 |
| Fig.3 ChIP-seq, Fig.4 RNA-seq (6hpf) | Murphy 2018 | GEO: GSE95033 |
| Fig.4 RNA-seq (0hpf, 3hpf) | Rougeot 2019 | GEO: GSE119070 |
| Fig.2, Fig.3, Fig4, CUT&Tag | This study<br>Akdogan-Ozdilek et al, 2022 | GEO: GSE237444, GSE178559 |
| Fig4. RNA-seq (12hpf) | Meng et al., 2022 | GEO: GSE197131 |
| **Experimental Models: Organisms/Strains** |  |  |
| Wild type zebrafish | Zebrafish International Resource Center | ZL57 |
| **Software and Algorithms** |  |  |
| HOMER | Heinz et al., Mol Cell, 2010 | http://homer.ucsd.edu/homer/motif/ |
| Bowtie2 | Langmead & Salzberg 2012 | https://www.nature.com/articles/nmeth.1923 |
| Samtools | Li et al., 2009 | https://samtools.sourceforge.net/ |
| Picard |  | http://broadinstitute.github.io/picard |
| g:Profiler | Raudvere et al., 2019 | https://academic.oup.com/nar/article/47/W1/W191/5486750 |
| rnaSTAR | Dobin et al., 2013 | https://academic.oup.com/bioinformatics/article/29/1/15/272537?login=true |
| bedtools | Quinlan and Hall 2012 | https://academic.oup.com/bioinformatics/article/26/6/841/244688 |
| Deeptools | Ramirez et al., 2014 | https://academic.oup.com/nar/article/42/W1/W187/2435511 |
| MACS2 | Zhang et al., 2012 | https://genomebiology.biomedcentral.com/articles/10.1186/gb-2008-9-9-r137 |
| Integrative Genomics Viewer (IGV) | Robinson et al., 2011 | https://www.nature.com/articles/nbt.1754 |
| R Packages, data visualization and parsing |  |  |
| dplyr | Wickham, 2023 | https://dplyr.tidyverse.org,<br>https://github.com/tidyverse/dplyr |
| pheatmap | Kolde, 2019 | https://CRAN.R-project.org/package=pheatmap |
| UMAP | McInnes, Healy, Melville, 2018 | https://arxiv.org/abs/1802.03426 |
| ggplot2 | Wickham, 2016 | https://ggplot2.tidyverse.org/ |

## Methods

### Zebrafish husbandry

Adult wild type Tübingen zebrafish were maintained on a 14h:10h light:dark cycle. Zebrafish husbandry and care were conducted in full accordance with animal care and use guidelines with ethical approval by the University Committee on Animal Resources at the University of Rochester Medical Center. Experimental samples were early zebrafish embryos ranged from 0hpf to 12hpf.

### Genomic profiling by CUT&Tag

6hpf and 12hpf H2A.Z and Anp32e genomic localization data were collected using Epicypher CUT&Tag protocol and library generation followed by Illumina paired-end sequencing. Briefly, embryos were collected and manually dechorionated, dissociated by pipetting and nuclei were extracted and collected. Around 100,000 nuclei were used for each replicate, Polyclonal H2A.Z antibody (Active motif, Cat. No 39113, 1:50 dilution) and Anp32e antibody (ThermoFisher Scientific Cat. No PA5-42860, 1:50 dilution) were used for CUT&Tag.

### Alignment, normalization, and peak-calling

For ChIP-seq, CUT&Tag, and ATAC-seq, fastq files were aligned to Zv11 using Bowtie2 (v2.2.5). Aligned .sam files were converted to .bam files using Picard sortsam. PCR duplicates were removed upon read count (RPKM) normalization and conversion to bigwig. This was done in one step using the deeptools application, bamCoverage. Peaks were called using MACS2 (v2.1.4) bdgpeakcall (-c 20 -g 100 -l 100) for ChIP-seq and ATAC-seq. All peak and promoter annotation set manipulations were performed using the bedtools (v2.30) suite. Bedtools intersect was used, where intersecting peaks are called with any amount of overlap. The ‘-v’ flag was used to derive non-intersecting intervals.

*Gene Ontologies (GO)*: GO analyses were performed using g:Profiler using default significance threshold. Bed files of any given feature set were inputted and associated with TSS of the nearest gene, matching genomic intervals to gene sets. Subsequently, functional enrichment analysis was performed.

### Accessibility analyses

Peaks were called on each individual ATAC-seq replicate before concatenating and merging any overlapping intervals between replicates. These peaks were first rank-normalized by sample and then variance sorted across genomic features, where features with the greatest variance in rank-normalized accessibility were selected. Upon clustering by accessibility trends across time, further analysis of these peaks was carried out in the absence of rank-normalization.

### Motif analyses

All motif analysis was performed using HOMER (v4.10). findMotifs.pl was used to identify enriched motifs within provided genomic features. Motifs identified in epiboly-specific putative CREs were defined against a background of dome-specific putative CREs. Motifs identified in Gastrulation accessible regions were defined against a union of all accessible regions comprising the timecourse. scanMotifGenomeWide.pl was used to query Zv11 for genomic intervals matching our selected Sox motif file (http://homer.ucsd.edu/homer/motif/HomerMotifDB/homerResults/motif335.motif). Towards our motif scan, sequence weights were maintained, but Log odds detection threshold was increased to 10 to generate a more highly specific motif set. Identified Sox motifs were intersected with a union of all accessible regions before further analysis.

### Transcriptomic analyses

All RNA-seq data were aligned and directly converted to duplicate-removed, RPKM-normalized .bam files using rnaSTAR (v2.7.8). Except where indicated, data were rank-normalized within samples to account for batch-to-batch variation and enable comparison across developmental time between studies.

### Plotting

Heatmaps and genomic averages were generated using deeptools (v3.5.1) plotHeatmap and plotProfile. All plots of promoters include promoter direction oriented left to right. R packages used for data parsing and visualization include dplyr, pheatmap, UMAP, and ggplot2. Boxplots were generated using standard R (v4.2.2). All figures were adjusted and polished in Affinity Designer.

### Statistical Methods

Students t-tests with R statistical software were used to determine statistical significance between boxplot means with a cutoff of P<=0.05 for all normally distributed data. Wilcoxon rank tests (non-parametric) were used for distributions that were non-normal. Data normality was evaluated using the Shapiro-Wilk test. Coding p-values for boxplot comparisons: *; **; *** for p<=0.05; p<=0.01; p<0.001.

## Data Availability

All sequencing datasets generated in this study were deposited to GEO data repository (GSE237444).

## Author contributions

F.N.H. and P.J.M designed experiments; F.N.H. and P.J.M performed experiments and data analyses; F.N.H., F.W.M., P.J.M prepared and wrote the manuscript.

## Acknowledgements

We would like to thank everyone in the Murphy lab for helpful discussions. This work was supported by the NIH (R35GM137833 to P.J.M) and De Kiewet fellowship (to F.N.H).

## Supplemental Figure 1

A. Enrichment averages of H3K4me1 and H3K4me3 (ChIP-seq) at blastula-specific, gastrula-specific, and shared peaks.

B. Boxplots of H3K27ac (ChIP-seq), accessibility (ATAC-seq), and 2A.Z (CUT&Tag) at gastrula-specific H3K4me1 regions. All three demonstrate increases from blastula to gastrula

C. Boxplots of H3K27ac, accessibility, and H2A.Z at gastrula-specific H3K4me3 regions. All three demonstrate increases from blastula to gastrula

D. List of GO terms for H3K4me3 gastrula-specific regions

E. List of GO terms for H3K4me1 gastrula-specific regions

## Supplemental Figure 2

A. Heatmap resultant from k-means and hierarchical clustering of rank-normalized accessibility scores

B. List of GO terms for Gastrula-specific and MZT-specific regions

## Supplemental Figure 3

A. Heatmaps of H2A.Z enrichment (CUT&Tag) at early gastrula and somitogensis across promoters

B. Heatmaps of Anp32e enrichment (CUT&Tag) at early gastrula and somitogensis across all promoters

C. H2A.Z enrichment (ChIP-seq) in WT and anp32e -/- embryos across all promoters Heatmaps of ANP32E enrichment

## References

1. Gibney & Nolan. Epigenetics and gene expression. Heredity 105, 4–13 (2010).

2. Pálfy, M., Schulze, G., Valen, E. & Vastenhouw, N. Chromatin accessibility established by Pou5f3, Sox19b and Nanog primes genes for activity during zebrafish genome activation. PLOS Genetics (2020).

3. Pijuan-Sala, B. et al. Single-cell chromatin accessibility maps reveal regulatory programs driving early mouse organogenesis. Nat Cell Biol. 22, 487–497 (2020).

4. Cusanovich, D. et al. The cis-regulatory dynamics of embryonic development as single-cell resolution. Nature 555, 538–542 (2018).

5. Giaimo, B., Ferrante, F., Herchenrother, A., Hake, S. & Borggrefe, T. The histone variant H2A.Z in gene regulation. Epigenetics and Chromatin.

6. Dijkwel, Y. & Tremethick, D. The Role of Histone Variant H2A.Z in Metazoan Development. Developmental Biology 10, (2022).

7. Brunelle, M. et al. The histone variant H2A.Z is an important regulator of enhancer activity. Nucleic Acids Research 43, 9742–9756 (2015).

8. Jin, C. et al. H3.3/H2A.Z double variant-containing nucleosomes mark ‘nucleosome-free regions’ of active promoters and other regulatory factors. Nature Genetics 41, 941–945 (2009).

9. Henikoff, S. Nucleosome destabilization in the epigenetics regulation of gene expression. Nature Reviews 9, 15–26 (2008).

10. Henikoff, S. & Smith, M. Histone Variants and Epigenetics. CSH Perspectives 7, (2015).

11. Subramanian, V., Fields, P. & Boyer, L. H2A.Z: a molecular rheostat for transcriptional control. Faculty Opinions (15AD).

12. Obri, A. et al. ANP32E is a histone chaperone that removes H2A.Z from chromatin. Nature 648–653 (2014).

13. Mao, Z. et al. Anp32e, a higher eukaryotic chaperone directs preferential recognition for H2A.Z. Cell Research 24, 389–399 (2014).

14. Murphy, P., Wu, S., James, C., Wike, C. & Cairns, B. Placeholder Nucleosomes Underlie Germline-to-Embryo DNA Methylation Reprogramming. Cell 993–1006 (2018).

15. Greenberg, R., Long, H., Swigut, T. & Wysocka, J. Single Amino Acid Change Underlies Distinct Roles of H2A.Z subtypes in Human Syndrome. Cell 178, 1421–1436 (2015).

16. Raja, D. et al. Histone variant dictates fate biasing of neural crest cells to melanocyte lineage. Development (2020).

17. Ruff, G., Murphy, K., Smith, Z., Vertino, P. & Murphy, P. Subtype-Independent ANP32E Reduction During Breast Cancer Progression in Accordance with Chromatin Relaxation. BMC Cancer 21, (2021).

18. Sarkar, A. & Hochedlinger, K. The Sox Family of Transcription Factors: Versatile Regulators of Stem and Progenitor Cell Fate. Cell Stem Cell 12, 15–30 (2013).

19. Schock, E. & LaBonne, C. Sorting Sox: Diverse Roles for Sox Transcription Factors During Neural Crest and Craniofacial Development. Frontiers Physiology 11, (2020).

20. Elizabeth Heeg-Truesdell & LaBonne, C. Wnt Signaling: A Shaggy Dogma Tale. Current Biology 16,.

21. Lukoseviciute, M. et al. From Pioneer to Repressor: Biomdal foxd3 Activity Dynamically Remodels Neural Crest Regulatory Landscape In Vivo.

22. Bogdanovic, O. et al. Dynamics of enhancer chromatin signatures mark the transition from pluripotency to cell specification during embryogenesis. Genome Research 2043–2053 (2012).

23. Kimmel, C., Ballard, W., Kimmel, S., Ullmann, B. & Schilling, T. Stages of embryonic development of the zebrafish. Developmental Dynamics 253–310 (1995).

24. White, R. et al. A high-resolution mRNA expression time course of embryonic development in zebrafish. eLife (2017).

25. Bannister, A. & Kouzarides, T. Regulation of Chromatin by Histone Modifications. Cell Research 21, 381–395 (2011).

26. Meng, F., Murphy, K., Makowski, C. & Murphy, P. Genome-wide chromatin accessibility is restricted by ANP32E. Nature Communications (2020).

27. Fuglerud, B. et al. A c-Myb mutant causes deregulated differentiation due to impaired histone binding and abrograted pioneer factor function. Nucleic Acids Research 45, 7681–7696 (2017).

28. Martik, M. & Bronner, M. Regulatory Logic Underlying Diversification of the Neural Crest. Trends Genet. 33, 715–727 (2017).

29. Herchenrother, A. et al. The H2A.Z and NuRD associated protein HMG20A controls early head and heart developmental transcription programs. Nature Communications 14, (2023).

30. Pünzeler, S. et al. Mutivalent binding of PWWP2A to H2A.Z regulates mitosis and neural crest differntiation. EMBO 36, 2263–2279 (2017).

31. Berta, D. et al. Deficient H2A.Z deposition is assoicated with genesis of uterine leiomyona. Nature 596, 398–403 (2021).

32. Xiong, Z., Wang, L., Wang, Q. & Yuan, Y. LncRNA MALAT1/miR-129 axis promotes glioma tumorigenesis by targeting SOX2. J Cell Mol Med 3929–3940 (2018).

