## Supplementary figures and images for "Anp32e protects against accumulation of H2A.Z at Sox motif containing promoters during zebrafish gastrulation"

### Supplementary Figures

Supplemental Figure 1

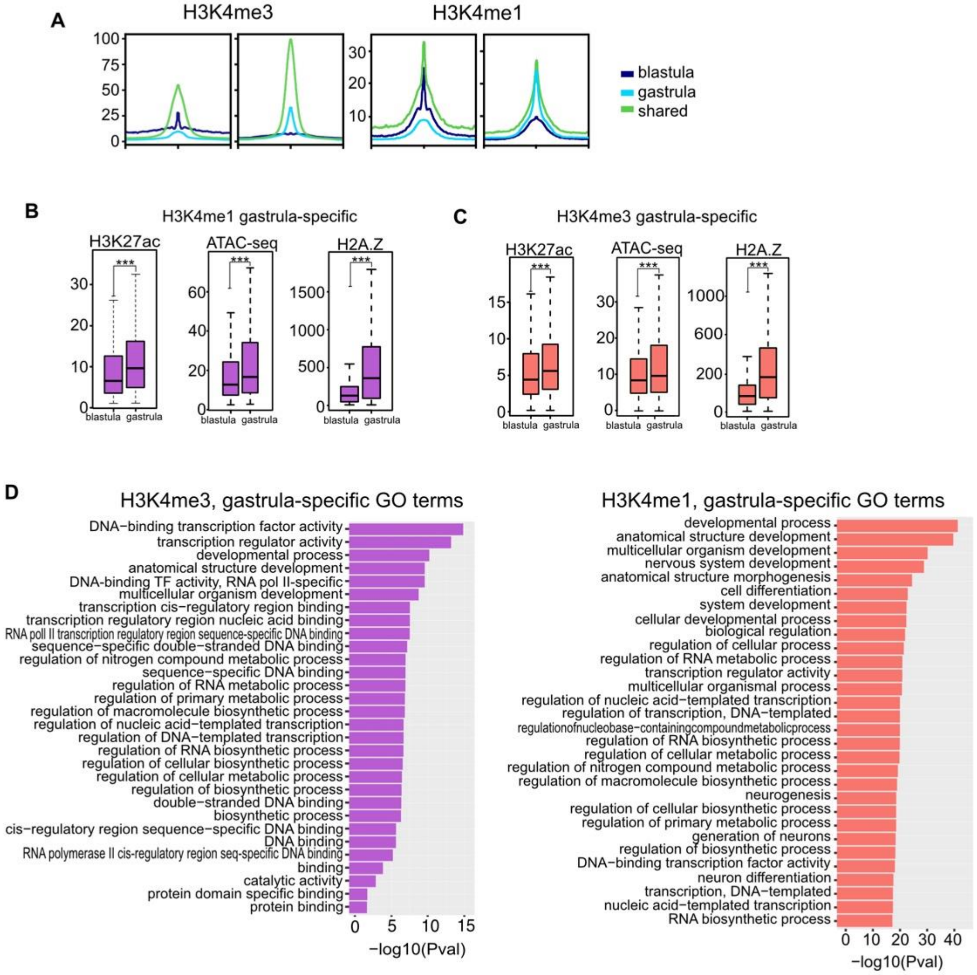

Supplemental Figure 2

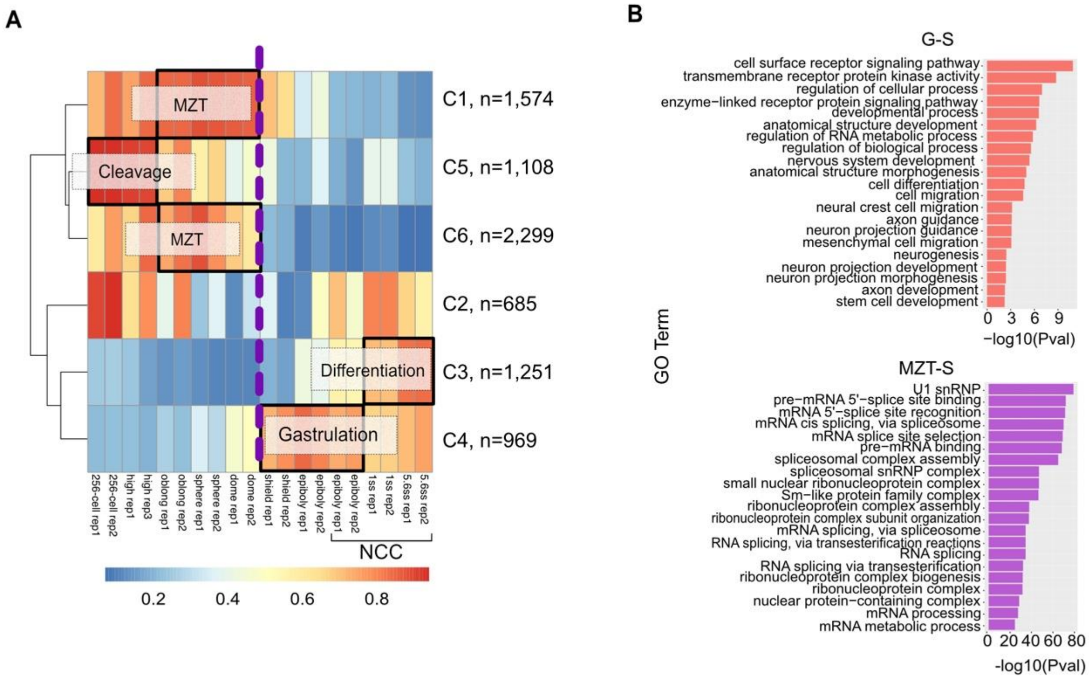

Supplemental Figure 3

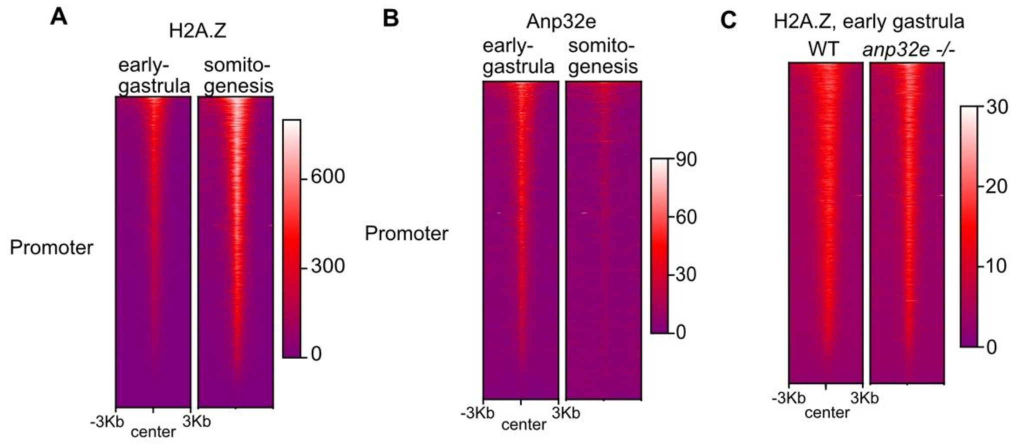
